# Fatty acid metabolism changes in association with neurobehavioral deficits in animal models of fetal alcohol spectrum disorders

**DOI:** 10.1101/2022.06.04.494839

**Authors:** Hye Mee Hwang, Yuka Imamura Kawasawa, Aiesha Basha, Shahid Mohammad, Kazue Hashimoto-Torii

## Abstract

Fetal alcohol spectrum disorders (FASD) show various behavioral problems due to prenatal alcohol exposure (PAE). Our previous study found significant changes in gene expressions linked to fatty acid metabolism in the brain of the PAE mouse model. Given the importance of fatty acids in normal brain functions and the contributions to neurodegenerative diseases, we hypothesized that the fatty acids changed by PAE contribute to neurobehavioral deficits in FASD. This study found an increase of palmitic acid and arachidonic acid in phospholipid compositions in the cerebral cortex of PAE at postnatal day 30. The increase of palmitic acid was consistent with the increase of the producing enzyme, fatty acid synthase (*Fasn*). The decrease of 26:6 fatty acid was also found in phospholipid. It is consistent with the increase of the Elongation of very long chain fatty acids protein 4 (ELOVL4) which uses 26:6 as a substrate for making very long chain fatty acids. However, there was no increase in the elongated products. Rather, we found an accumulation of the lipid droplets (LDs) in the PAE brain, suggesting changes in fatty acid metabolism that lead to the accumulation of excessive fatty acids. Although metabolic measurements, including plasma triglyceride level, were not affected by PAE, the abundance of fatty acid-related gut microbiota was altered. Interestingly, multi-omics association analysis revealed a potential contribution of the altered gut microbiota, primarily *Ruminococcaceae* that produces short chain fatty acid, to LD formation in the PAE brain and the behavioral problems, suggesting that the gut microbiome could serve as a tool to facilitate uncovering the brain pathophysiology of FASD and a potential target to mitigate neurobehavioral problems.

## Introduction

Alcohol exposure during gestation adversely affects fetal development. Fetal alcohol spectrum disorder (FASD) is an umbrella term describing a group of clinical conditions resulting from prenatal alcohol exposure (PAE). Those conditions include facial dysmorphology and cognitive and neurobehavioral deficits (Doney et al., 2014; Heck et al., 2008; Streissguth et al., 1990; Williams et al., 2015; Wozniak et al., 2019). Recent clinical and preclinical animal studies demonstrated that PAE is associated with metabolic dysregulation after birth, including increased body fat, a higher incidence of obesity, and hypertension (Cook et al., 2019; Fuglestad et al., 2014; Z. He et al., 2015; Weeks et al., 2020), highlighting the metabolically vulnerable individuals with PAE. Of note, metabolic disruption was shown to associate negatively with cognition in adolescents (Meo et al., 2019) and positively with the risk of developing dementia such as Alzheimer’s disease (AD) in adults (Gustafson et al., 2003). Disrupted lipid homeostasis was reported in the brain biopsy of other neurodegenerative diseases, including amyotrophic lateral sclerosis and Parkinson’s diseases (PD) (Brekk et al., 2020; Derk et al., 2018; Dupuis et al., 2008). In these patients, accumulation of lipid droplets (LDs) was found in various brain regions and cell types (Brekk et al., 2020; Derk et al., 2018; Paula-Barbosa et al., 1980). LDs are organelles that store excess fatty acids to protect cells from lipid toxicity (Cohen, 2018), and the biogenesis and degradation of LDs are tightly coupled with cellular metabolism to maintain homeostatic lipid levels (Rambold et al., 2015).

Alcohol interacts directly with fatty acid to produce fatty acid ethyl ester, thereby identified as a biomarker of maternal alcohol drinking (Himes et al., 2015). Furthermore, in pregnant mothers who drink alcohol, fatty acid composition in their plasma was different between mothers with offspring showing abnormal development and those showing normal development (Sowell et al., 2020). In animals exposed to alcohol throughout gestation, docosahexaenoic acid (DHA), an n-3 fatty acid, was reduced in phospholipid collected from the postnatal hippocampus (Kim, 2008; Wen & Kim, 2004). In another study, administration of DHA between P11 and P20 improved social behavior deficits of PAE rats (Wellmann et al., 2015). In FASD young children, choline that enhances the fatty acid oxidation in the liver (Li et al., 2018) was supplied for 9 months, and improved non-verbal IQ and working memory in those children (Wozniak et al., 2020). These studies collectively indicate that disturbed fatty acid metabolism can be the treatment target for FASD.

Gut microbiota crucially regulates host metabolism (Vuong et al., 2017) and modulates brain function via the gut-brain axis (Carabotti et al., 2015). Short chain fatty acids (SCFAs) produced by gut microbiota are one of the major metabolites that influence host lipid metabolism, and accumulating evidence demonstrates that the SCFA also affects neurobehavior through the gut-brain axis (Dalile et al., 2019; Silva et al., 2020). A number of studies also found differential abundances of gut microbial compositions in the feces of patients who suffer from neurodegenerative diseases (Fang et al., 2020). Animal studies also demonstrated that changing gut microbial composition affects neuropathology and behavior (Kim et al., 2020; Sun et al., 2018).

Similar to the adult cases described above, several studies have reported that both maternal and early postnatal microbiota are essential for exploratory, social, and sensorimotor behavior development in mice (Degroote et al., 2016; Hsiao et al., 2013). Depleting maternal microbiota by antibiotic treatment impaired thalamocortical axonogenesis of the fetus, and the replenishment of the *Clostridia* bacteria into the mother by oral gavage improved the impairment (Vuong et al., 2020). In FASD, there are a few recent studies reporting changes in the gut microbiome in animal models of FASD (Bodnar et al., 2022; Virdee et al., 2021; Wang et al., 2021). However, how these changes are associated with the neurobehavioral phenotypes remains unknown.

Using a mouse model of PAE, this study revealed for the first time that learning deficits and anxiety phenotype are associated with the disturbance of fatty acid metabolism in the brain and gut microbiota, but not with other metabolic measurements such as body weights, plasma triglycerides, or blood glucose levels. These results suggest that the gut microbiome may serve as a sensitive biomarker for lipid dysregulation and have crucial contributions to lipid-mediated brain pathology and neurobehavioral issues, irrespective of blood metabolic molecules in FASD.

## Results

### PAE increases the expression of a fatty acid synthesizing enzyme in the frontal cortex at a juvenile stage

In our previous study, single cell RNA sequencing revealed that the expressions of genes in the fatty acid biosynthesis pathway, such as elongation of very long chain fatty acids protein 4 (*Elovl4*) and fatty acid synthase (*Fasn*), were significantly upregulated in the cortical neurons in the mouse primary motor cortex at P30, long after acute prenatal alcohol exposure once a day at 4.0g/kg weight at embryonic day 16 and 17 (Mohammad et al., 2020). ELOVL4 is a fatty acid elongase required to form long chain fatty acids greater than 28 carbon lengths long (Agbaga et al., 2008). To examine whether the protein level is also increased, the number of ELOVL4-positive cells was compared between PAE and the control brains (offspring of PBS received dams) by immunohistochemistry at P30 when both motor skill learning deficits and anxiety were observed in PAE mice (Hwang & Hashimoto-Torii, 2022; Mohammad et al., 2020).

Consistent with the increase at the RNA level (Mohammad et al., 2020), the number of cells that express ELOVL4 proteins was significantly increased in the PAE primary motor cortex (Figure 1A, B). We also found that a nearby cortical region, the cingulate cortex that is involved in controlling anxiety (Hwang & Hashimoto-Torii, 2022; McClure et al., 2007; Monk et al., 2008), shows an increase in the number of ELOVL4-positive cells in PAE mice (Figure 1 – figure supplement 1), suggesting the increase of ELOVL4 in the frontal cortex of PAE mice. Consistent with the previous mouse study in which ELOVL4 protein expression was examined in normal adult mouse brains (Sherry et al., 2017), most of the ELOVL4 immunolabeling was found in the soma of cells that express NeuN, a marker of mature neurons, in control animals (Figure 1C). In addition, the intracellular distribution pattern of ELOVL4 in the neurons was not altered by PAE (Figure 1B, Figure 1 – figure supplement 1B, D). Given that PAE did not alter the number of ELOVL4 expressing cells in the cerebellum (Figure 1 – figure supplement 1C) that also controls motor learning (de Zeeuw & ten Brinke, 2015), fatty acid composition and/or metabolism are anticipated to be altered uniquely in neurons of the frontal cortex of PAE mice.

**Figure 1.**
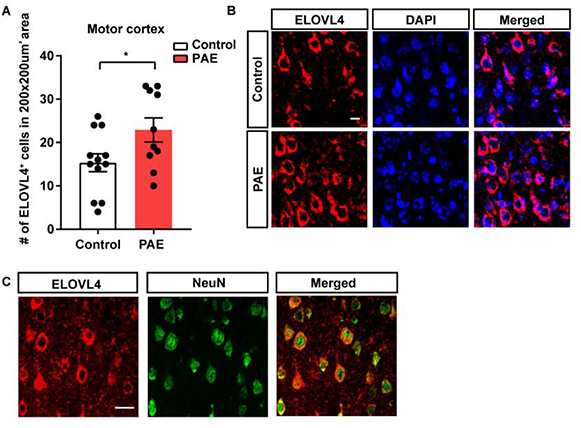
PAE increases neuronal ELOVL4 expression in the motor cortex at P30. (A) The number of ELOVL4-positive cells is significantly higher in the motor cortex of PAE mice compared to the control mice (p=0.038). *p<0.05. Student’s t-test. Control n=12, PAE n=10. (B) Representative images for immunolabeling of ELOVL4 (red) and DAPI (blue). Scale bar = 10µm. (C) Representative images of coronal section from control cortex demonstrate that ELOVL4 expression (red) is detected in NeuN-positive neurons (green). Scale bar = 25µm. Graphs represent mean ± SEM. Each dot represents an individual animal. **Figure 1 – source data 1** **Quantification of the number of ELOVL4 positive cells in the motor cortex.**

### The fatty acid composition of phosphatidylethanolamine in the cell membrane is altered in PAE motor cortex at the juvenile stage

We then examined the fatty acid compositions in the cell membrane in the motor cortex at P30 by analyzing fatty acid in phospholipid using liquid chromatography tandem mass spectrometry. Phosphatidylcholine (PC) and phosphatidylethanolamine (PE) are the most abundant phospholipids in cell membranes (van der Veen et al., 2017). PC is found mostly in the outer layer of the plasma membrane, whereas PE is found in the inner layer of the plasma membrane and mitochondrial membrane (van der Veen et al., 2017). Therefore, we analyzed the fatty acids in both PC and PE.

First, we examined total amounts of PC and PE and found no significant changes in PAE mice compared to control mice (Figure 2 – figure supplement 1A, B). Disturbance of the PC/PE ratio is known to alter energy metabolism and is associated with non-alcoholic fatty liver diseases and obesity (van der Veen et al., 2017). Therefore, we examined the PC/PE ratio. Similar to another study (Choi et al., 2018), the PC/PE ratio indicated a higher amount of PE than PC in cortices of both control and PAE groups. However, their ratio was not altered by PAE (Figure 2 – figure supplement 1C).

In the comparisons of each fatty acid molecular species, a significant increase was found only in the palmitic acid (16:0) and the arachidonic acid (20:4) in PE of PAE mice (Figure 2A). Palmitic and arachidonic acids are saturated fatty acids (SFA) and polyunsaturated fatty acids (PUFA), respectively, both of which are highly abundant in the cell membrane (Bazinet & Layé, 2014; Carta et al., 2017). The increase in palmitic acid was consistent with an increase in *Fasn* RNA expression (Mohammad et al., 2020) as the function of the FASN is to synthesize palmitic acid (Maier et al., 2006). In PC, there was no significant difference in any of the fatty acid molecular species between PAE and control mice (Figure 2 – figure supplement 2A).

**Figure 2.**
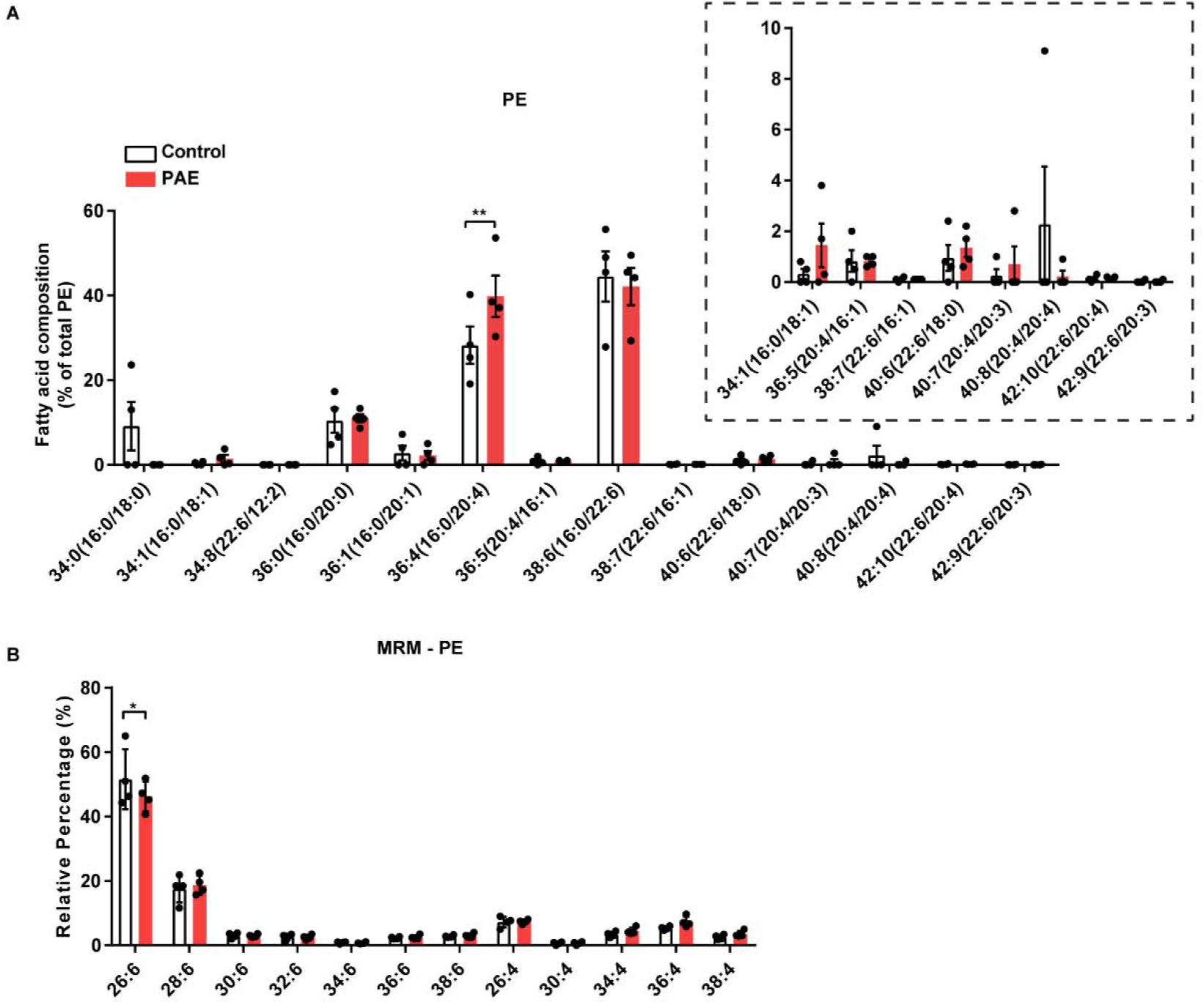
Fatty acid composition in PE is altered by PAE. (A) PAE mice show a significant increase in abundance of 36:4 of PE (16:0/20:4) compared to control mice (p=0.0096) in the motor cortex at P30. Inset shows the magnified view of the fatty acid species that are less abundant. (B) Multi reaction monitoring (MRM) analysis reveals a significant decrease of 26:6 of PE fatty acid in PAE motor cortex compared to control (p=0.0397). No interaction between fatty acid species and prenatal exposure types by two-way ANOVA. Bonferroni’s multiple comparisons tests were used as post hoc tests. Control (PBS exposure) n=4, PAE n=4. *,**p<0.05, 0.01. Graphs represent mean ± SEM. Each dot represents an individual animal. PE = phosphatidylethanolamine. **Figure 2 – source data 1** **Analyzed relative percentage of fatty acid composition in PE of the motor cortex.**

The Multi reaction monitoring (MRM) was used to detect very long chain fatty acids that ELOVL4 specifically produces in both PC and PE. ELOVL4 is involved in elongating both SFA and PUFA by adding 2 additional carbons to synthesize very long chain fatty acids that are longer than 28 carbons (C28); thus, any of the fatty acids that are in between C26 and C36 are thought to be substrates for ELOVL4 (Agbaga et al., 2008; Hopiavuori et al., 2019). In the retina, PUFAs elongated by ELOVL4 are incorporated into PC (Deák et al., 2019). Therefore, we measured PUFA between C26 and C36 in PC but also in PE, in which we observed changes in fatty acids in PAE cortex (Figure 2B, and Figure 2 – figure supplement 2B). The analysis revealed that only 26:6 PUFA, the shortest fatty acid substrate of ELOVL4 enzyme, was significantly decreased in PE of PAE cortex (Figure 2B). However, no difference was observed in any of the fatty acid molecules of PC between control and PAE mice (Figure 2 – figure supplement 2B). The reduction of ELOVL4 substrates was consistent with the increase of ELOVL4 in the cortex of PAE mice. However, unexpectedly, the increase of the very long chain fatty acids that ELOVL4 produce was not observed. This indicated a possibility that those very long chain fatty acids are metabolized immediately after the production by excessive ELOVL4.

### Accumulation of lipid droplets (LDs) in PAE brain

The results described above suggested that ELOVL4-produced very long chain fatty acids are metabolized in PAE motor cortex, or potentially sequestered away to prevent lipid toxicity. LDs store such intracellular fatty acids by incorporating them into neutral lipids such as triglycerides, while excessive accumulation of LDs is one of the pathological signatures in neurodegenerative diseases and aging brains (Olzmann & Carvalho, 2019). In addition, an increase in both palmitic acid and arachidonic acid, which showed a significant increase in PAE motor cortex (Figure 2A), promotes the formation of LDs in hepatic cells and monocytes (Fujimura & Usuki, 2012; Guijas et al., 2012). Altogether, we hypothesized that PAE might facilitate formation of LDs in the motor cortex.

To detect LDs, we used oil red o (ORO), which stains neutral triglycerides and lipids in the brain. We first confirmed ORO staining with a 12-month-old mouse as LDs accumulate in aging brains (Shimabukuro et al., 2016). ORO-labeled LDs were found inside cells not only in the motor cortex but also in cingulate and piriform cortices, striatum, hippocampus, and lateral ventricle (LV) wall where periventricular glial cells are located (Figure 3 – figure supplement 1), similar to observations made in aging studies (Hamilton et al., 2015; Shimabukuro et al., 2016).

We then examined LD accumulation in the motor cortex and several brain regions of PAE and control mice at P30, as depicted in Figure 3A. The number of LD accumulating cells was significantly increased in the motor cortex of PAE mice compared with that of control mice (Figure 3B, C). Notably, a significant increase of LD included cells was also observed in other brain regions such as the striatum, CA3, and dentate gyrus of the hippocampus (Figure 3B). Although statistically not significant, the piriform and cingulate cortices and LV wall showed trends of increase in the number of cells with ORO labeling. LDs were immunolabelled by using antibodies against their surface proteins, such as perilipin (PLIN) 1,2 and 3 (Marschallinger et al., 2020; Shimabukuro et al., 2016). Using double immunostaining of PLIN1 and NeuN, we found that the LDs accumulate in the neurons in PAE mice.

**Figure 3.**
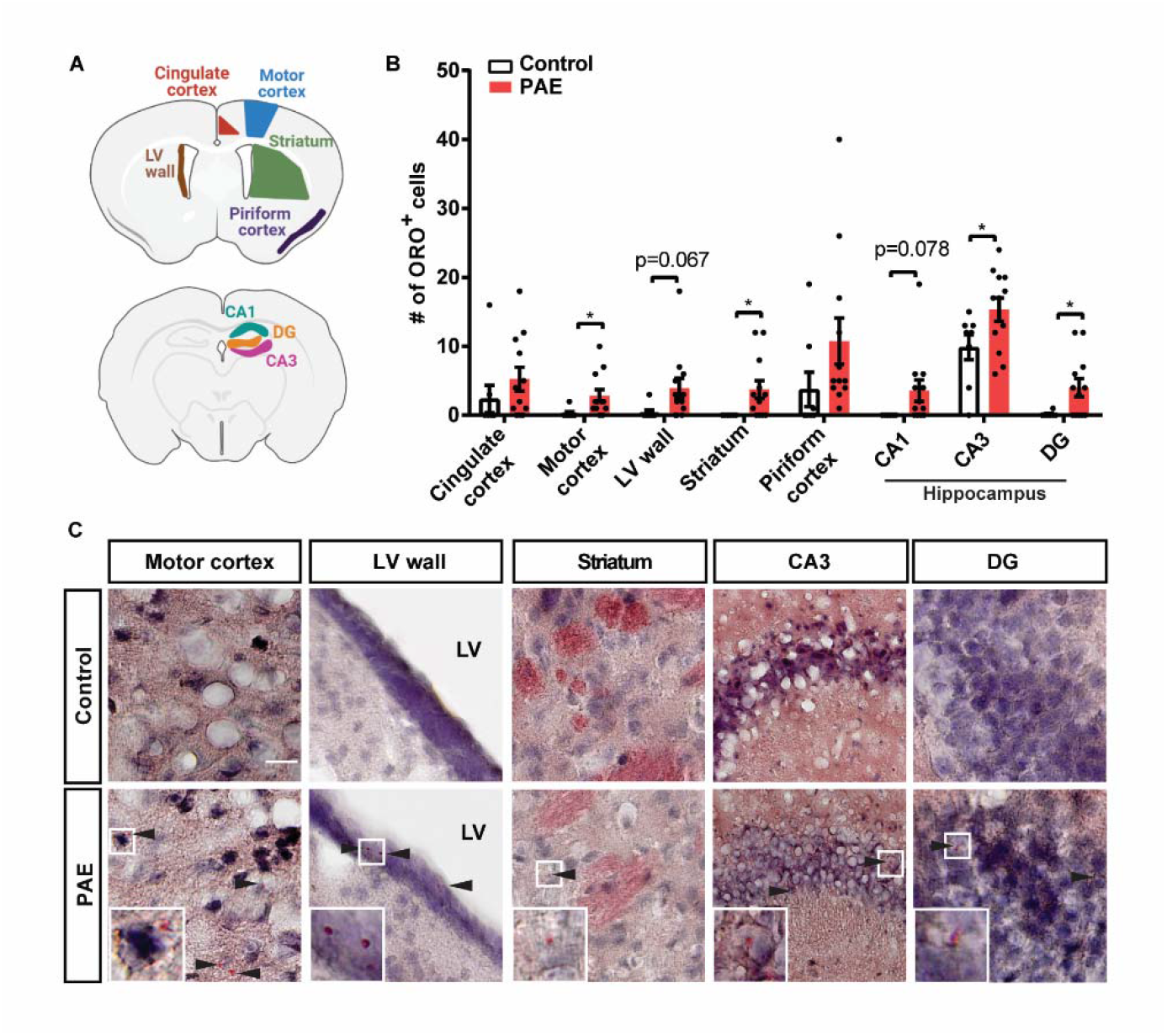
LDs are accumulated in various brain regions in PAE mice at P30. (A) Highlighted areas in the schema indicate brain regions analyzed in mice. (B) The numbers of LD included cells were quantified in 200 x 200 µm^2^ area in each brain region. The data show an increase of LD accumulating cells in the motor cortex (p=0.0352), striatum (p=0.0311), both DG and CA3 of the hippocampus (p=0.0267 and 0.0466, respectively), and lateral ventricle (LV) wall (p=0.067) in PAE mice. *p<0.05. Student’s t-test. Control n=8; PAE n=12. Graphs represent mean ± SEM. Each dot represents an individual animal. (C) Representative images of ORO stained brain areas show significant changes in the number of cells accumulating LDs. Insets show higher magnification views of the ORO-positive cells. Scale bar=20µm. Arrowheads indicate LDs. **Figure 3 – source data 1** **Quantification of the number of LD accumulated cells in various brain regions.**

Next, we examined the LD droplet formation during postnatal development. Given that the dam’s milk contains a high level of fat at approximately 30% (Görs et al., 2009) and high fat induces LD accumulation in periventricular glial cells adjacent to LV wall in mice (Ogrodnik et al., 2019), there was a possibility that the LD formation may be complicated around the weaning when the pups are no longer suckling. Thus, the number of ORO-positive cells in the brain was examined on three postnatal days, before weaning (P15), the day of weaning (P20), and after weaning (P25).

We found that the LV wall and the two regions of gray matter in the cortex showed different dynamics of LD depositions during normal development (Figure 3 – figure supplement 2). In PAE, the dynamics were also different between the LV wall and the gray matter (Figure 3 – figure supplement 2). Only at P20 (and P30 in Figure 3B), but not at P15 or P25, the differential accumulations of LDs were observed between PAE and control cortices (Figure 3 – figure supplement 2), suggesting spatiotemporal dynamics of LD formation around weaning date.

### The gut microbiome that are associated with fatty acid biogenesis and metabolism are altered by PAE

Accumulation of LDs is one of the neuropathological hallmarks in neurodegenerative diseases such as PD and AD. These patients also show changes in both peripheral triglycerides levels (Bernath et al., 2020; Fu et al., 2020) and gut microbiota composition (Petrov et al., 2017; Zhuang et al., 2018), suggesting ingrained changes in lipid metabolism in these patients. Given that our PAE mice accumulated LDs in the brain and previous studies showed an association between PAE and metabolic disorders such as obesity and hyperlipidemia (Cook et al., 2019; Fuglestad et al., 2014; Weeks et al., 2020), we examined body weight, plasma triglyceride level, and blood glucose level in our animals. However, none of those metabolic measurements were altered in our acute PAE mouse model (Figure 4 – figure supplement 1), indicating neurobehavioral problems without manifest metabolic dysfunctions in PAE.

Then we examined if the gut microbiota composition is altered by PAE. The gut microbiota produces SCFAs such as butyrate, acetate, and propionate. SCFAs regulate the biogenesis, oxidation, and metabolisms of fatty acids in various tissues, likely in context dependent manner (J. He et al., 2020). Some types of SCFAs inhibit the functions of a histone deacetylase to improve memory and learning in normal animals (Intlekofer et al., 2013; Vecsey et al., 2007) and an animal model of meningitis (Barichello et al., 2015), while others lead to autism-like behavior issues (Macfabe et al., 2007). This interplay of the gut microbiome and neuronal signaling, called gut-brain axis, is mediated by bidirectional molecular mediators, including bioactive lipids that can modulate the gut-brain axis (Baptista et al., 2020). Therefore, we examined an association between the changes in neurobehavioral phenotype and gut microbiota.

As shown in the experimental timeline in Figure 4A, following collections of fecal pellets for 16S ribosomal RNA (rRNA) sequencing, animals were placed on an accelerated rotarod to assess their motor learning by conducting 3 trials per day for two consecutive days. The next day, animals were placed in an elevated plus maze (EPM) to assess anxiety. The behavioral test results showed that PAE mice have both motor learning deficit and anxiety phenotype that were demonstrated by a significant reduction in the learning index and time spent in the open arm, respectively (Figure 4B, C, D). In addition to reduced open arm time in EPM, the numbers of open arm entries and closed arm entries, as well as the total number of entries in arms were decreased in PAE mice (Figure 4 – figure supplement 2A,B,C). However, there was no significant difference in the time spent in closed arms between control and PAE mice (Figure 4 – figure supplement 2D), while PAE mice spent significantly more time in the center (Figure 4 – figure supplement 2E). The increased center time but a decreased open arm time in PAE mice indicated that the mice showed exploring behavior but avoid entering the open arm. Moderate but significant correlations between motor learning index and anxiety measurement, open arm time in EPM, were also found (Figure 4 – figure supplement 3).

**Figure 4.**
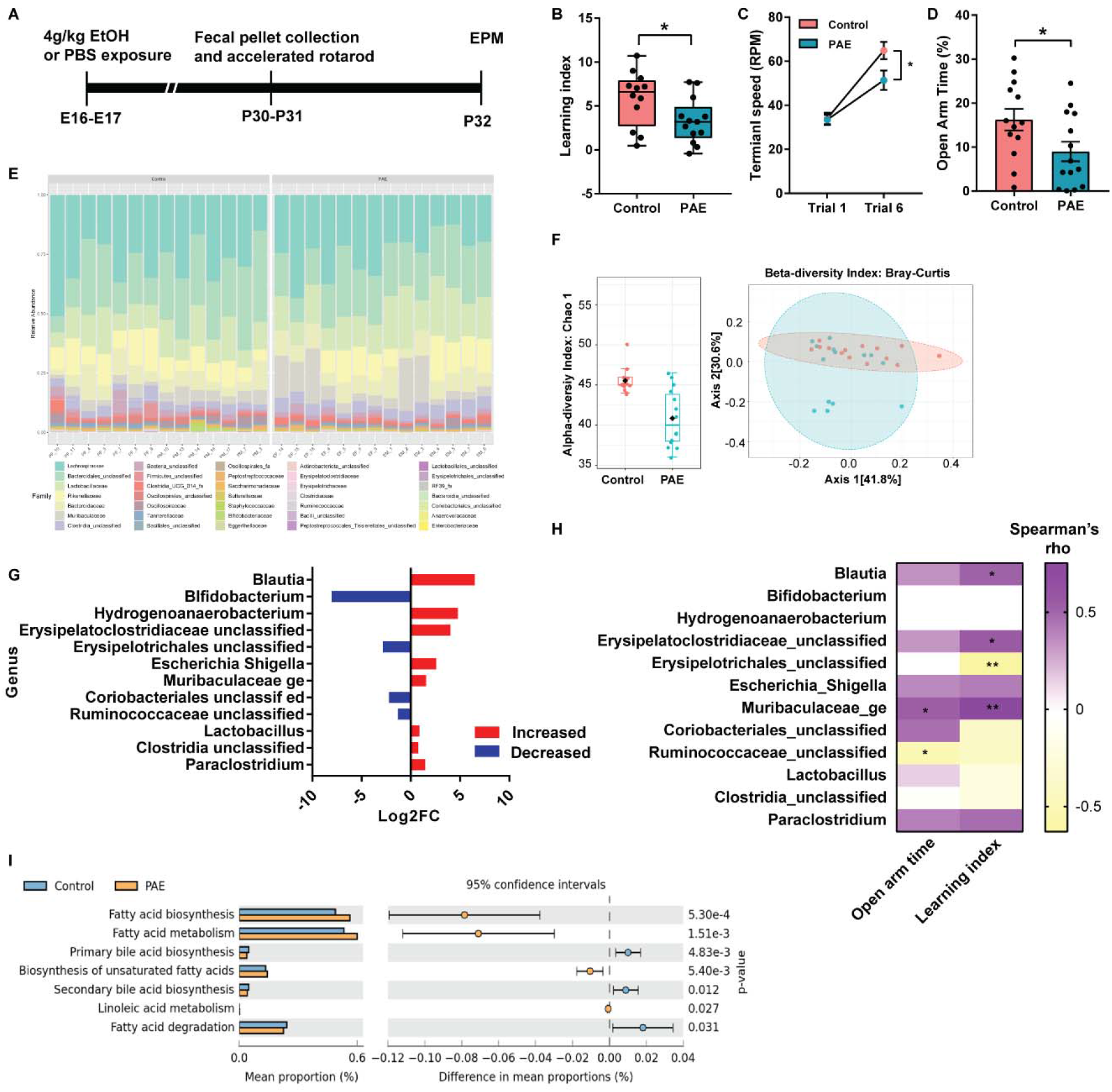
Gut dysbiosis is associated with behavioral problems in PAE mice. (A) Timeline of the experiment. (B) The PAE mice score significantly lower in the learning index in the accelerated rotarod test (p=0.0351). Student’s t-test. Control n=12; PAE n=13. Graph represents a box blot with 25th, median, and 75th percentiles, and whiskers extend to the lowest and highest values. Each dot represents an individual animal. (C) PAE mice reach significantly lower terminal speed in the last trial (Trial 6) compared to control mice (adj.p=0.0257). Two-way ANOVA revealed no significant interaction between trials and exposure. Tukey’s post hoc test was used for multiple comparisons. Control n=12; PAE n=13. Each dot on the graph represents a group mean ± SEM. (D) PAE mice spend significantly less time in the open arm compared to the control mice (p=0.0370). Student’s t-test. Control n=13; PAE n=14. (C,D) Graph represents mean ± SEM. Each dot represents an individual animal. (E) Stacked bar plots show the taxonomic composition of the microbial community at the family level in each animal used for analysis. (F) Microbial diversity analysis reveals that PAE mice have significantly smaller alpha diversity compared to that of control mice (p=0.00030926) as well as differential microbial communities (beta diversity). Alpha diversity was compared using Chao1 at the genus level. Graph represents a boxplot with 25th, median, and 75th percentiles. Whiskers represent 9 and 91 percentiles, and outside of whiskers represent outliers. Each dot represents an individual animal, and the black diamond represents the mean. Beta diversity analysis at the genus level shows that PAE and control mice have differential microbial communities. F = 3.2946, R^2^ = 0.11246, p=0.017 by Permutational ANOVA (PERMANOVA). Graph represents 2-D Principal Coordinate Analysis (PCoA). Each dot represents an individual animal. (G) Using EdgeR, 12 differentially abundant taxonomy are identified at the genus level. Graph represents log2 Fold Change (FC), and bacteria is listed in the order of the most significant (*Blautia*) to the least significant (*Paraclostridium*). Red bar = increased in PAE, Blue bar = decreased in PAE. (H) The abundance of bacterial taxa at the genus level in PAE mice correlates with levels of motor learning and anxiety. Color of heatmap indicates rho values by Spearman’s correlation analysis. (I) Functional profiles of the microbiome predict changes in fatty acid-related pathways in PAE mice. Graph represents the difference in mean proportions (%) with 95% confidence intervals calculated by Welch’s inverted method. *,**p<0.05, 0.01. **Figure 4 – source data 1** **Analyzed accelerated rotarod learning index, the terminal speed at trials 1 and 6, open arm time in EPM test, and Spearman’s rho and p-values to generate the heatmap.**

The 16s rRNA sequencing data was analyzed using Mothur (Schloss et al., 2009). After alignment and mapping with the SILVA v138 reference database, the operational taxonomic unit (OTUs) were clustered at 97% identity threshold to sort reads into each gut bacteria. OTUs with less than 4 read counts were removed from each sample, and then bacteria with its OTU counts showing less than 10 % prevalence in entire samples were removed from the analysis. After those filtrations, the final numbers of the total read counts were similar between samples, ranging from 81765 to 82281 (Figure 4 – figure supplement 4A). The plateau of rarefaction curves showed that most of the abundant species are included in each sample (Figure 4 – figure supplement 4B). The curves of PAE samples indicated higher variability in the number of detected OTUs (Figure 4 – figure supplement 4B).

As a result, 35 bacterial families were found in the samples; *Lachnospiraceae*, *Unclassified Bacteroidales,* and *Lactobacillaceae* were some of the highly abundant microbiota in the samples (Figure 4E). The bacterial diversity analysis revealed that PAE mice have significantly lower alpha diversity compared with control mice (Figure 4F). Furthermore, beta diversity based on Bray-Curtis distance matrix indicated a significant difference between control and PAE mice in microbial composition (Figure 4F).

We further investigated differentially abundant microbiota at the genus level. Analysis revealed that 12 bacterial genera (8 increased and 4 decreased in PAE) were differentially abundant between PAE and control (Figure 4G). To test the correlation between behavioral phenotypes and abundance of microbiota, we performed Spearman’s correlation analysis between the abundance of significantly altered bacterial genera and animal’s learning index in the accelerated rotarod test and time spent in the open arm of EPM in PAE group. The analysis revealed that the abundance of three microbiota, *Blautia, Unclassified Erysipelatoclostridiaceae,* and *Muribaculaceae_ge,* showed a positive correlation with the motor learning index, whereas *Unclassified Erysipelotrichales* showed a negative correlation with the index (Figure 4H). *Blautia* is a gut microbial genus that produces SCFAs such as butyric acid and acetic acid (Liu et al., 2015). *Muribaculaceae*, also known as *S24-7*, is a family of bacteria that produces the enzymes for carbohydrate degradation (Lagkouvardos et al., 2019). *Erysipelatoclostridiaceae* is a family of *Firmicutes* that are involved in carbohydrate metabolism (Ottman et al., 2012). With time spent in the open arm in EPM, *Muribaculaceae_ge* and *Unclassified Ruminococcaceae* showed positive and negative correlations, respectively (Figure 4H). *Ruminococcaceae* degrades cellulose and hemicellulose and subsequently converts these compounds to SCFAs, which can be absorbed and used for generation of energy by the host (Valentine et al., 2020).

Using TaxFun2, changes in functional profiles due to altered microbiota compositions were examined (Wemheuer et al., 2020). The data revealed that lipid metabolism is the most affected biological function by PAE (Figure 4 – figure supplement 5). We further examined which lipid metabolism-related pathways are altered in PAE mice and found that biosynthesis and metabolism of fatty acids and biosynthesis of unsaturated fatty acids are increased. Metabolism of linoleic acid, a PUFA that is a substrate to be elongated to the arachidonic acid (Whelan & Fritsche, 2013), was also increased (Figure 4I). On the other hand, fatty acid degradation was decreased in PAE mice (Figure 4I). Notably, the analysis also revealed that both primary and secondary bile acid biosynthesis is significantly decreased in PAE mice (Figure 4I). Bile acids help breaking down fats into fatty acids to facilitate their absorption into cells (McMillin & DeMorrow, 2016). The liver synthesizes primary bile acids from which secondary bile acids are transformed by gut bacteria (Chiang, 2013). Collectively, these results suggested that the fatty acid biosynthesis is increased while degradation is decreased in PAE mice, resulting in excessive depositions of fatty acids by the gut microbiota in PAE mice.

### *Ruminococcaceae* show the strongest correlation with LD accumulation in the brain and behavioral phenotypes

To comprehensively examine the correlations between obtained biological measures, we integrated data using a mixOmics-based R package called DIABLO (Rohart et al., 2017; Singh et al., 2019). Different from Spearman’s correlation analysis above, which used only PAE samples (Figure 4H), DIABLO algorithm enables the integration of all PAE and control samples to identify key features across different data types while discriminating between multiple phenotypic groups. The neurobehavioral phenotypes, brain LD, and microbiome results obtained from an animal in either PAE or control group were used for the analysis. As shown in Figure 5, the motor learning index showed a strong negative correlation with the total brain LD accumulations. Among the brain regions, the piriform cortex was one of the brain regions we saw a strong correlation with anxiety measures. The piriform cortex is mainly known for the odor processing region, but it also receives input from the basolateral amygdala, a brain region important for anxiety, to form a cortical circuit to shape responses to the threatening stimuli (East et al., 2021).

**Figure 5.**
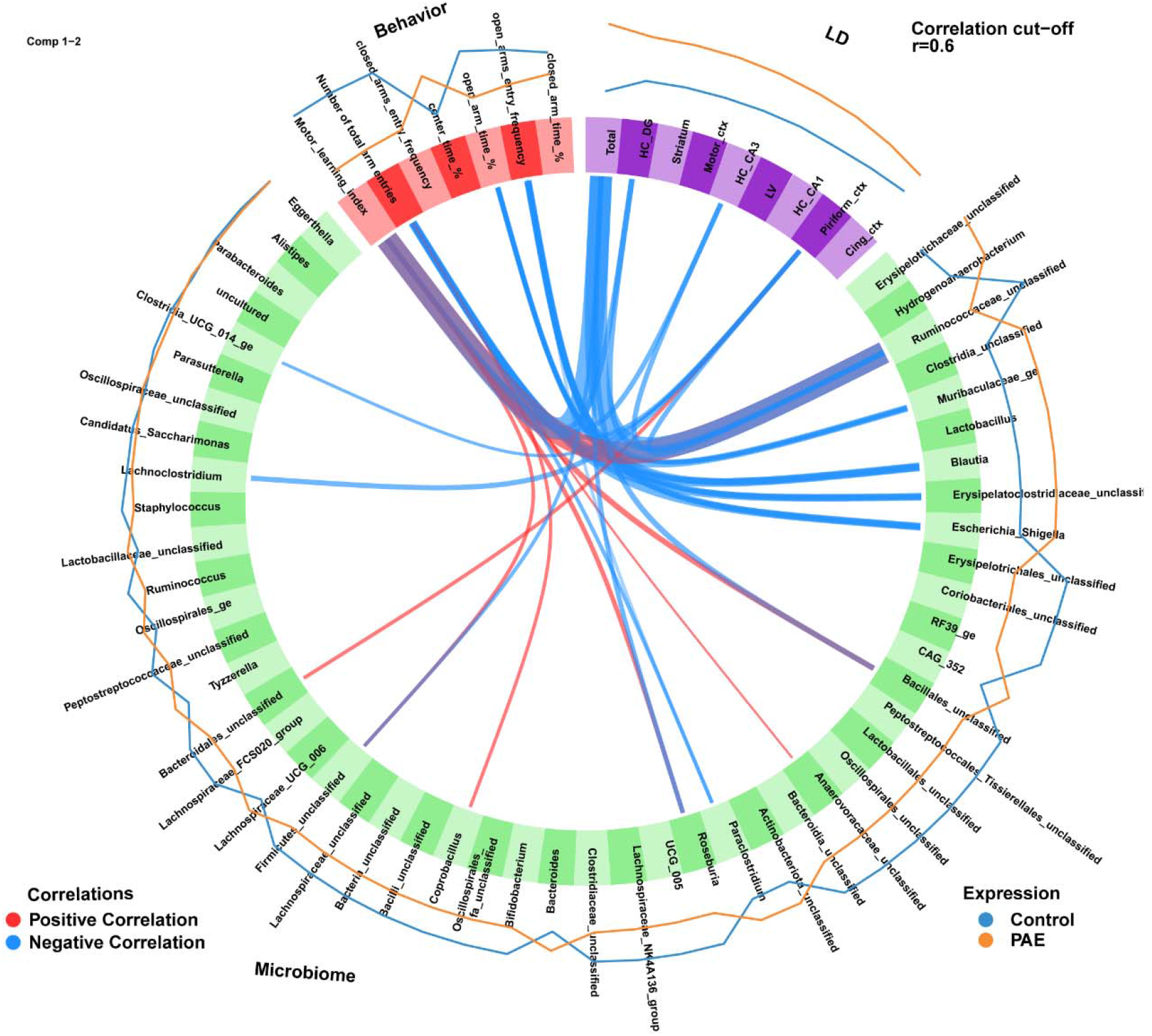
*Ruminococcaceae* is strongly correlated with brain LD accumulation and behavioral phenotype. Circos plot shows positive and negative correlations between the measures obtained in behavioral tests (accelerated rotarod and EPM), number of LD accumulating cells in the brain, and microbial abundance. The thickness of the link corresponds to the correlation coefficient (r). Control n=8, PAE n=12. HC = hippocampus, DG = dentate gyrus, LV = lateral ventricle wall.

There were stronger correlations with LD accumulation and neurobehavioral measures in the microbiota that are known to be associated with fatty acid synthesis and metabolisms, such as *Unclassified Ruminococcaceae* (Valentine et al., 2020)*, Blautia* (Liu et al., 2015), as well as carbohydrate metabolism associated microbiota including *Muribaculaceae_ge* (Lagkouvardos et al., 2019) and *Unclassified Erysipelatoclostridiaceae* (Ottman et al., 2012). In humans, consuming *Escherichia Shigella* contaminated food or water can cause serious illness (Zaidi & Estrada-García, 2014). Similarly, in mice, intraperitoneal administration of *Shigella* bacteria induced severe diarrhea and acute inflammation, similar to humans (Yang et al., 2014). In addition, *Escherichia Shigella* was shown to be increased in anxiety disorder patients (Jiang et al., 2018). Among all detected bacteria, *Unclassified Ruminococcaceae* showed the strongest negative correlation with the total number of accumulated LDs, but a positive correlation with the motor learning index, suggesting that *Ruminococcaceae* may affect the behavior through the impact on the lipid metabolism in the brain.

## Discussion

Increased expression of proteins encoding enzymes that are involved in fatty acid biogenesis and modification in the cortical neurons in PAE mice long after prenatal exposure (Figure 1A and previously published work (Mohammad et al., 2020)) led us to investigate the fatty acid composition in membrane phospholipid in the cortex (Figure 2, Figure 2 – figure supplement 2). We also applied MRM to quantify the very long chain fatty acid species in PAE and FASD research for the first time. Despite the increased expression of ELOVL4 in the cortex (Figure 1A), an enzyme that elongates the C26, there were no changes in C28 and the longer chain fatty acids (Figure 2B). However, the reduction of C26:6 in PE (Figure 2B) suggested a possibility that, despite increased usage of C26:6 by ELOVL4, the produced very long chain fatty acids are broken down immediately after the elongation; therefore, no significant increase was observed in phospholipid.

We also found an increase in palmitic and arachidonic acids in PE in PAE mice (Figure 2A). Of note, previous studies showed an increase of arachidonic acids in maternal plasma from mothers who consumed alcohol during pregnancy and had children with lower cognitive test scores compared to those mothers who consumed alcohol but had children with the normal scores (Sowell et al., 2020). In the same study, palmitic acids in maternal plasma were also positively correlated with alcohol intake.

No changes in the amount of the fatty acids that are longer than C26 and simultaneous reduction of C26 in the plasma membrane of the PAE brain cells indicated a possibility that excessively elongated very long chain fatty acids are being metabolized and sequestered by forming LDs to prevent lipidopathy. In fact, we observed the increase of LDs in various brain regions in PAE mice (Figure 3). Another possible explanation as to why there was no increase of C26 observed despite the increase of ELOVL4 in the PAE cortex might be due to the specific subcellular localization of these very long chain fatty acids: such as synaptic vesicles (Hopiavuori et al., 2018) while we collected PC and PE of all membrane fractions that potentially mask the differences in the synaptic compartment.

Our study revealed for the first time the accumulation of the LDs in the brains of PAE animals (Figure 3) that show various cognitive and behavioral problems (Hwang & Hashimoto-Torii, 2022; Mohammad et al., 2020). There were slight differences in the levels of accumulation between brain regions; however, all of the brain regions showed an increase of LDs in PAE, suggesting brain-wide issues. The variability in LD accumulation levels could be due to the difference in lipid composition among different brain regions, where the prefrontal cortex and motor cortex show similar lipid composition in the membrane but are different from other brain regions such as the hippocampus, striatum, and cerebellum (Fitzner et al., 2020). In addition, behavioral measurements also showed correlations with the LD accumulations in the brain regions that are known to be involved in motor learning and anxiety (Figure 5). Interestingly, the LDs appeared as transient pathological features that are likely to be affected by diet during postnatal development. In our observation, the accumulation of LDs was dynamic during the weaning time (Figure 3 – figure supplement 2).

In the functional annotation to changes in the microbial profiles in PAE animals, we found an increase in the fatty acid synthesis pathway but a reduction in fatty acid degradation, suggesting a possible systemic increase in amounts of fatty acids in PAE mice (Figure 4I). This prediction was consistent with the lipid-related brain pathology and changes in fatty acid contents in membrane phospholipids in the brain (Figure 2). Further functional analysis of microbiome revealed that the metabolic pathway of linoleic acid, which is a precursor fatty acid for arachidonic acid (Whelan & Fritsche, 2013), was also increased in PAE mice (Figure 4I). This change might be associated with the increase of the arachidonic acid in PAE brains (Figure 2A). The primary and secondary bile acid synthesis were also predicted to be changed in PAE’s microbiome profiles (Figure 4I). This finding was interesting because the changes in bile acid biogenesis were observed from the biosignature of microbiome-derived metabolites in both alcohol-exposed dams and their fetuses (Virdee et al., 2021). Collectively, these biosignatures predicted from changes in gut microbial compositions suggest that the systemic changes in fatty acid biogenesis and metabolism and that those may be linked to the changes in PAE brains.

Association analysis between neurobehavior, lipid-associated brain pathology, and gut microbiome revealed the strong correlations of microbiota that are involved in fatty-acid biogenesis and metabolism with brain pathology and the behavior (Figure 5). Among those microorganisms, *Unclassified Ruminococcaceae* showed the strongest associations with both brain pathology and behavior. *Ruminococcaceae* are found in both human and mouse gut microbiome, but the abundance is higher in humans than in mice (Nguyen et al., 2015). Interestingly, the abundance of *Ruminococcaceae* in the gut is associated in both positive and negative directions with neurobehavioral measures, depending on which health measures and which brain functions were used (Renson et al., 2020; Sun et al., 2019; Vogt et al., 2017). For example, *Ruminococcaceae* was associated with lower anger, greater cognitive functions, and varied personality traits in humans (Renson et al., 2020). In addition, AD patients showed a reduced abundance of *Ruminococcaceae* (Vogt et al., 2017). On the other hand, autistic patients showed higher abundance compared with the control subjects (Sun et al., 2019). Furthermore, hepatic encephalopathy patients treated with an oral capsular fecal microbial transplant that includes enriched *Ruminococcaceae* showed improvement in cognitive problems (Bajaj et al., 2019). Therefore, *Ruminococcaceae* may serve as a biomarker or/and become a probiotic treatment option for the FASD. The integrative analysis (Figure 5) also showed an intriguing possibility that the gut microbiota might directly affect the lipid pathogenesis without changing metabolism in the entire body at a significant level. An exciting hypothetical mechanism is that metabolites of gut microbiota affect vagus nerve signaling (Fülling et al., 2019) and that vagus nerve stimulation directly affects lipid composition in various brain regions, including the striatum and motor cortex (Surowka et al., 2015), which could be mediated by the gut-brain axis (Baptista et al., 2020).

Although changes in systemic metabolic measures such as blood glucose level and body weight have been reported in chronic alcohol exposure animal models (Chen & Nyomba, 2003; Yao & Grégoire Nyomba, 2007) and adult FASD patients (Weeks et al., 2020), those measures were not altered by acute prenatal alcohol exposure in our animal model (Figure 4 – figure supplement 1), suggesting no relation with neurobehavioral problems or lipid-associated brain pathology. This may be due to milder phenotypes after acute PAE compared to a chronic PAE model. However, importantly, our model exhibits various neurobehavioral problems, including motor learning deficits and anxiety, and disruption in fatty acid related metabolism in brain and gut. Therefore, our results suggest that the effects of gut microbiota may profoundly and directly affect the brain pathology and function in PAE.

## Materials and Methods

### Animals

All animal experiments were conducted in accordance with the National Institutes of Health Guide for the Care and Use of Laboratory Animals and approved by the Institutional Animal Care and Use Committee at Children’s National Hospital (Protocol no. 00030323). To expose pregnant dam to alcohol, 4.0 g/kg body weight of ethanol (PAE group) or PBS (control group) was injected intraperitoneally (i.p.) at E16 and E17 to timed pregnant CD-1 (strain code: 022) mice that were purchased from Charles River Laboratories (Wilmington, MA) as previously done (Mohammad et al., 2020). The dams were randomly assigned to PAE or control group without any predetermined criteria. The day of birth was designated as postnatal day (P) 0, and female and male mice around the age of P30 were used for all experiments, otherwise specified in the figure legends. All the animals were maintained on a light-dark cycle (lights on 6:00–18:00) at a constant temperature (22 ± 1°C).

### Accelerated rotarod

The test was performed as previously described (Mohammad et al., 2020). Briefly, the mice were placed on a rotating bar, and the length of time that they could retain their balance during acceleration of rotation to a max speed of 80 rpm in 5 min was recorded. The testing phase consisted of 2 consecutive days with three trials per day. Each trial was at least 15 minutes apart and was terminated when the mouse fell off, made one complete rotation without walking on the rotating rod, or reached maximum speed after the 5-min session. The motor learning index was calculated by averaging the difference in terminal speed of the two consecutive trials.

### Elevated plus maze test (EPM)

The maze is a grey plus-shaped apparatus with two open arms and two closed arms linked by a central platform. Mice were individually put in the center of the maze facing an open arm and allowed to explore the maze for 300 seconds. A video was recorded during the experiment and analyzed with MouBeat ImageJ Plugin as per the user guide as previously done (Hwang & Hashimoto-Torii, 2022).

### Immunohistochemistry

Mice were deeply anesthetized with isoflurane (Henry Schein, Melville, NY) and perfused transcardially with 10 ml of ice-cold PBS followed by 10 ml of chilled 4% paraformaldehyde (PFA). The brains were removed and immerse-fixed in 4% PFA at 4 °C overnight. Then incubated in 10% and 30% sucrose in PBS for 24 hr sequentially at 4 °C and embedded in the OCT compound (cat# 4583; VWR, Randor, PA). Coronal sections were cut at 20 or 50µm on a cryostat (CM3050S; Leica, Buffalo Grove, IL).

Free-floating staining was performed with 50µm thick brain sections. Briefly, antigen retrieval was performed following the manufacturer’s protocol (cat# 00-4955-58; ThermoFisher, Waltham, MA) when necessary. Then sections were incubated in hydrogen peroxide in methanol (1:4) solution for 30 min at −20°C to inactivate endogenous peroxidase activity. After rinsing with PBS containing 0.01% of Tween-20 (PBS-T), sections were incubated with 2% BSA for 30 min at room temperature for blocking. Sections were then incubated with ELOVL4 (1:500 gifted from Dr. Robert Anderson or 1:200; cat# 224608; Abcam, Waltham, MA), and/or NeuN (1:300; cat# MAB3777; EMD Millipore, Burlington, MA) primary antibodies for overnight at 4 °C. For ELOVL4 immunolabeling, 2hr incubation with biotinylated anti-rabbit IgG (cat# 711-065-152; Jackson ImmunoResearch, West Grove, PA) diluted at 1:300 followed by 1hr incubation with avidin-biotin complex (1:1:100; cat# 32020; ThermoFisher) and 1hr incubation with TSA plus Cyanine-3 (1:300; cat# NEL744001KT; Akoya Biosciences, Marlborough, MA) was performed. For NeuN staining, sections were incubated for 2hr with HRP-conjugated anti-mouse IgG diluted at 1:300 and followed by 1 hr incubation of TSA plus Cyanine-2 (1:300; cat#NEL745001KT, Akoya Biosciences). Sections were counterstained with DAPI and mounted on slides with CC/Mount mounting medium (cat# C9368; Sigma, St. Louis, MO). Images were acquired using a confocal microscope (FV1000; Olympus Center Valley, PA). All images were analyzed with ImageJ using the Cell Counter tool.

### Oil Red O (ORO) staining

20µm thick brain sections were prepared as described above. For staining, ORO stock solution was prepared by dissolving 0.05g of ORO powder (cat#O0625; Sigma) in 10mL of isopropanol. Then 60% ORO working solution was prepared freshly with distilled water and filtered before use. Brain slices were incubated in 60% ORO solution for 10 min, washed thoroughly with distilled water, and incubated for 15 min with hematoxylin for counterstaining. Then the slices were mounted on slides with CC/Mount mounting medium, and images were acquired with an Olympus VS120 microscope. Brightness and contrast were adjusted with CellSens, and ORO-positive cells were manually counted with ImageJ Cell Counter tool.

### Lipidomics of Phospholipid Fatty Acids

Motor cortical regions were dissected from both hemispheres of P30 mice and snap-frozen in liquid nitrogen. Samples were stored at −80°C and shipped to Lipid Analysis Core at Emory University for phospholipid fatty acid quantification. Lipids were extracted from the cortical tissues using Bligh and Dyer method (Bligh & Dyer, 1959), and extracted lipids were directly loaded onto the mass spectrometer (SCIENX QTRAP LC-MS/MS system; Framingham, MA) for targeted lipidomics. The phospholipid class was selectively targeted using characteristic scans, and the same mass spectrometer was operated in multi reaction monitoring (MRM) mode to detect specific species of very long chain fatty acids that are presumably synthesized by ELOVL4 (Matraszek-Zuchowska et al., 2016). Detected fatty acids are converted to relative percentages by dividing the peak intensity of each fatty acid species by the total peak intensities detected per animal. To compare total PC and PE contents, all peak intensities from an individual animal were summed and divided by the total weight of the tissue.

### Plasma collection and measurement of triglycerides

Whole blood was collected from non-fasted and overnight fasted (16 hr) mice in an EDTA-treated tube (cat# 365974; BD, Franklin Lakes, NJ) by cardiac puncture with a 23-25G needle from a deeply anesthetized animal. Cells were removed from the plasma by centrifugation at 2000 x g at 4°C for 10 min. The supernatant was carefully transferred to a clean centrifuge tube by pipetting and stored in a −80°C freezer until further analysis. Measurement of triglycerides was carried out by a commercially available kit (cat# ab65336; Abcam) that detects glycerol levels after converting triglycerides to free fatty acids and glycerol by following the manufacturer’s protocol. Absorbance was measured with xMARK microplate spectrophotometer (cat# 1681150; Bio-rad, Hercules, CA).

### Fecal pellet collection and the 16S rRNA sequencing

Fecal pellets were collected from P30 animals and put on dry ice immediately after collection. DNA was extracted using the MagMAXTM CORE Nucleic Acid Purification Kit (Thermo Fisher Scientific, Waltham, MA), according to manufacturer instructions, and subjected to the library preparation for 16s rRNA V3-V4 region sequencing by using repliQa HiFi ToughMix (Quantabio, Beverly, MA). Next-generation sequencing was performed on an Illumina MiSeq by 2 x 300 bp paired-end readings. Mothur (Schloss et al., 2009) was used to align and map sequences with the SILVA v138 database. Chimeric reads were removed with VSEARCH, and operational taxonomic units (OTU) were clustered at 97% similarity. Biom files generated by Mothur were put into MicrobiomeAnalyst (https://www.microbiomeanalyst.ca/) for downstream analysis after removing low counts less than 4 and prevalence in samples less than 10%. Filtered data were scaled by total sum scaling, and alpha and beta diversity were determined by Chao1 and Bray-Curtis Index, respectively. Differential abundance analysis was carried out using EdgeR. To perform functional profiles analysis, representative OTU FASTA sequences and OTU count tables were retrieved from Mothur and analyzed with “Tax4fun2” package in R (Wemheuer et al., 2020). Statistical Analysis of taxonomic and functional profiles (STAMP) (Parks et al., 2014) was used to perform statistical analysis to compare metagenomic functional profiles between control and PAE mice.

### Multi-omics integrative correlation analysis

DIABLO multi-omics integration method from the mixOmics R package (Rohart et al., 2017) was applied to integrate different types of datasets from the same animals that have measurements in all three data sets: microbiome, data of the numbers of cells with lipid droplet (LD) in the brain regions, and results of behavioral tests (accelerated rotarod and EPM tests). We found 8 control and 12 PAE mice that meet this criterion. Centered log ratio (Clr) transformed microbiome and Z-score transformed the number of LDs accumulated in the brain, and behavioral measurements were used as the input data. The data integration was carried out following the manual. Briefly, a matrix model was designed to connect all datasets with a link strength at 1 between two components were used to generate the final DIABLO model. The circos plot shows the correlation strength and directionality between variables of different types of our datasets with a correlation coefficient cut off > |0.6|.

### Statistical analysis

All histological data were acquired from the defined subregions of the brain. All groups consisted of mice from at least two different litters with a similar number of female and male mice otherwise noted in the figure legends. The number of n indicated in the figure legends is that of biological replicates. The sample size for each experiment was determined based on previous experience in similar experiments. For all in vivo experiments, no animals were excluded from the analysis. Fatty acid analysis (Figure 2 and Figure 2 – figure supplement 2) and 16s RNA sequencing (Figure 4) were analyzed by the personnel who were group-blinded. Behavioral experiments were performed unblinded; however, automated analysis was used for EPM analysis (Figure 4). All the immunohistological analysis was done group-blinded by the investigator. Plasma triglycerides, blood glucose, and body weight measurements were performed unblinded (Figure 4 – figure supplement 1). All of the statistical analysis was carried out with GraphPad Prism 7.01. We performed the D’Agostino–Pearson to test the normality of data. For the data that passed the normality test, Student’s t-tests or one-way or two-way ANOVA was used. Post hoc Tukey’s or Bonferroni’s test was done as described in the figure legends. Simple main effects were reported when there was a statistically significant interaction between independent variables by two-way ANOVA. Pearson’s or Spearman’s correlation coefficient calculation was done for normally or non-normally distributed data, respectively. P values of less than 0.05 were considered statistically significant.

### Data availability

All original 16s rRNA sequencing raw data haven been deposited to the National Library of Medicine Bioproject under accession ID PRJNA842719. Data generated or analyzed in this study are included in the manuscript and Source Data 1 file.

## Acknowledgments

We would like to thank Dr. Robert E. Anderson for kindly providing ELOVL4 primary antibody. This study was supported by F31AA027693 (H.M.H), R01AA025215, R01AA026272 (K.H-T), and 1U54HD090257-01 from the NIH, District of Columbia Intellectual and Developmental Disabilities Research Center Award (DC-IDDRC) program. The 16S rRNA sequencing work was processed at the Genome Sciences and Bioinformatics Facility at the Penn State University College of Medicine. The lipid analysis was supported in part by the Emory Integrated Lipidomics Core (EILC), which is subsidized by the Emory University School of Medicine and is one of the Emory Integrated Core Facilities. Additional support was provided by the Georgia Clinical & Translational Science Alliance of the National Institutes of Health under Award Number UL1TR002378. The content is solely the responsibility of the authors and does not necessarily reflect the official views of the National Institutes of Health.

---

**Figure 1 – figure supplement 1.**
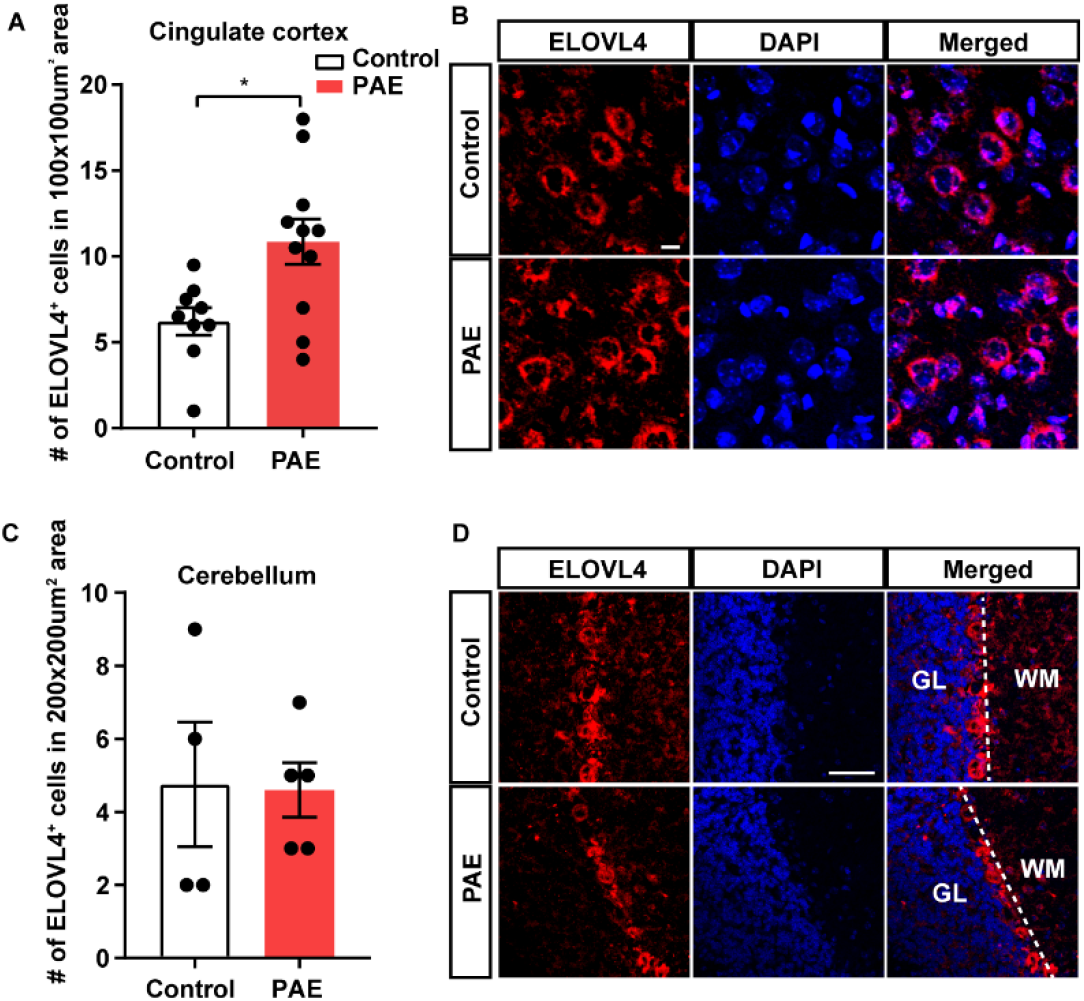
The number of ELOVL4 expressing cells is increased in the cingulate cortex but not in the cerebellum. (A) The number of ELOVL4-positive cells is significantly higher in the cingulate cortex of PAE mice compared with control mice at P30 (p=0.011). *p<0.05. Student’s t-test. Control n=9, PAE n=11. (B) Representative images of ELOVL4 (red) and DAPI (blue) staining in the cingulate cortex. Scale bar = 10µm. (C) The number of ELOVL4-positive cells in the cerebellum is not different between control and PAE mice at P30. Control n=4, PAE n=5. (D) Representative images of ELOVL4 immunostaining in the cerebellum. Scale bar = 50 µm. Graphs represent mean ± SEM. Each dot represents an individual animal. GL = granular layer, WM = white matter. **Figure 1 – figure supplement 1 – source data 1** **Quantification of the number of ELOVL4 positive cells in the cingulate cortex and cerebellum.**

**Figure 2 – figure supplement 1.**
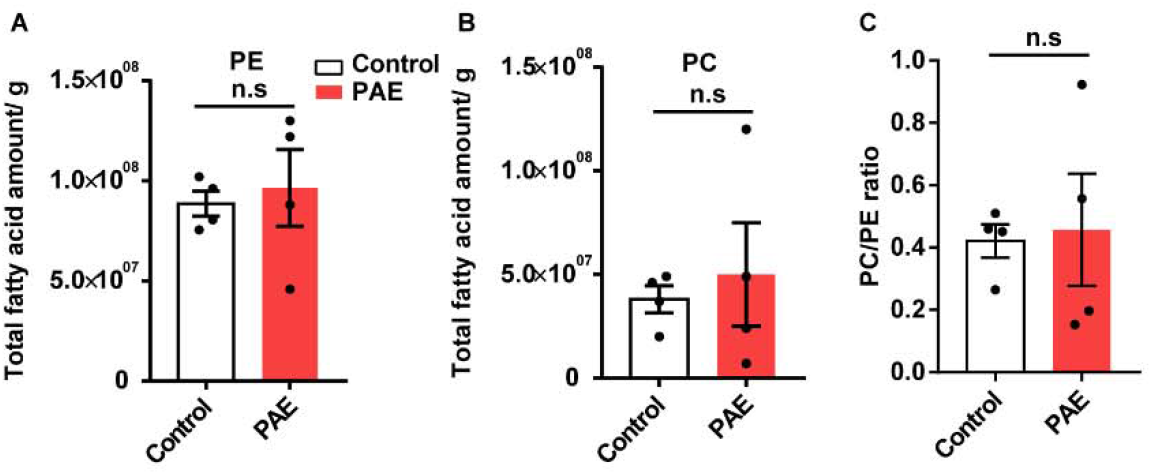
PAE does not alter the total amount of PC or PE from the motor cortex at P30. (A-C) There is no significant difference in the total amount of PC (A) and PE (B) or the ratio of PC/PE (C) between PAE and control animals by Student’s t-test. Control n=4, PAE n=4. Graphs represent mean ± SEM. Each dot represents an individual animal. PE = phosphatidylethanolamine, PC = phosphatidylcholine, n.s = not significant. **Figure 2 – figure supplement 1 – source data 1** **Quantification of the total fatty acid amount of PC and PE and their ratio.**

**Figure 2 – figure supplement 2.**
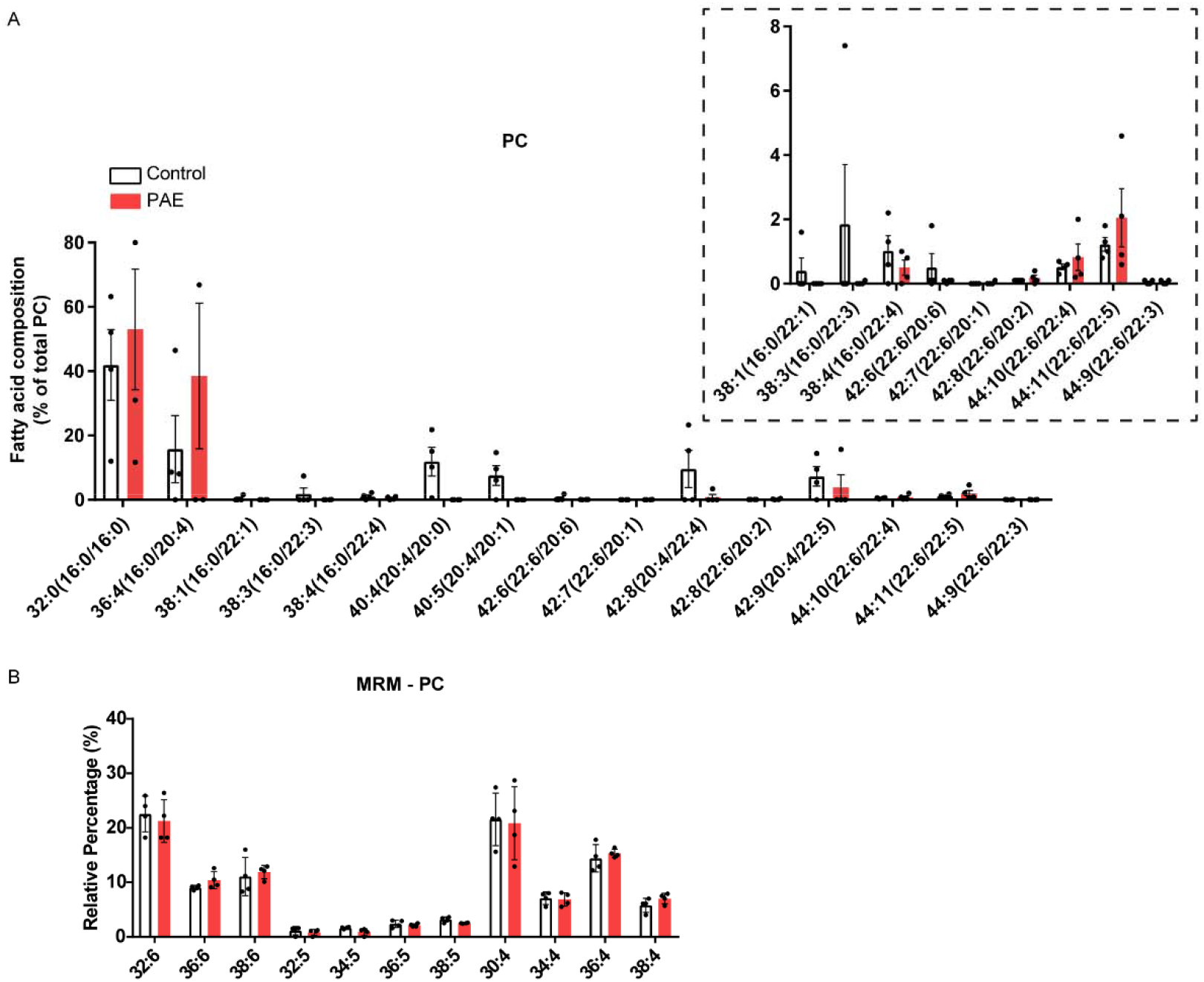
PC fatty acid composition in the motor cortex is not affected by PAE. (A, B) There is no difference in abundance of fatty acid species between control and PAE in PC of the motor cortex at P30. Two-way ANOVA found no interaction between fatty acid species and exposure types. Control n=4, PAE n=4. Inset shows the magnified view of the fatty acid species that are less abundant. Graphs represent mean ± SEM. Each dot represents an individual animal. PC = phosphatidylcholine, MRM = multi reaction monitoring. **Figure 2 – figure supplement 2 – source data 1** **Analyzed relative percentage of fatty acid composition in PC of the motor cortex.**

**Figure 3 – figure supplement 1.**
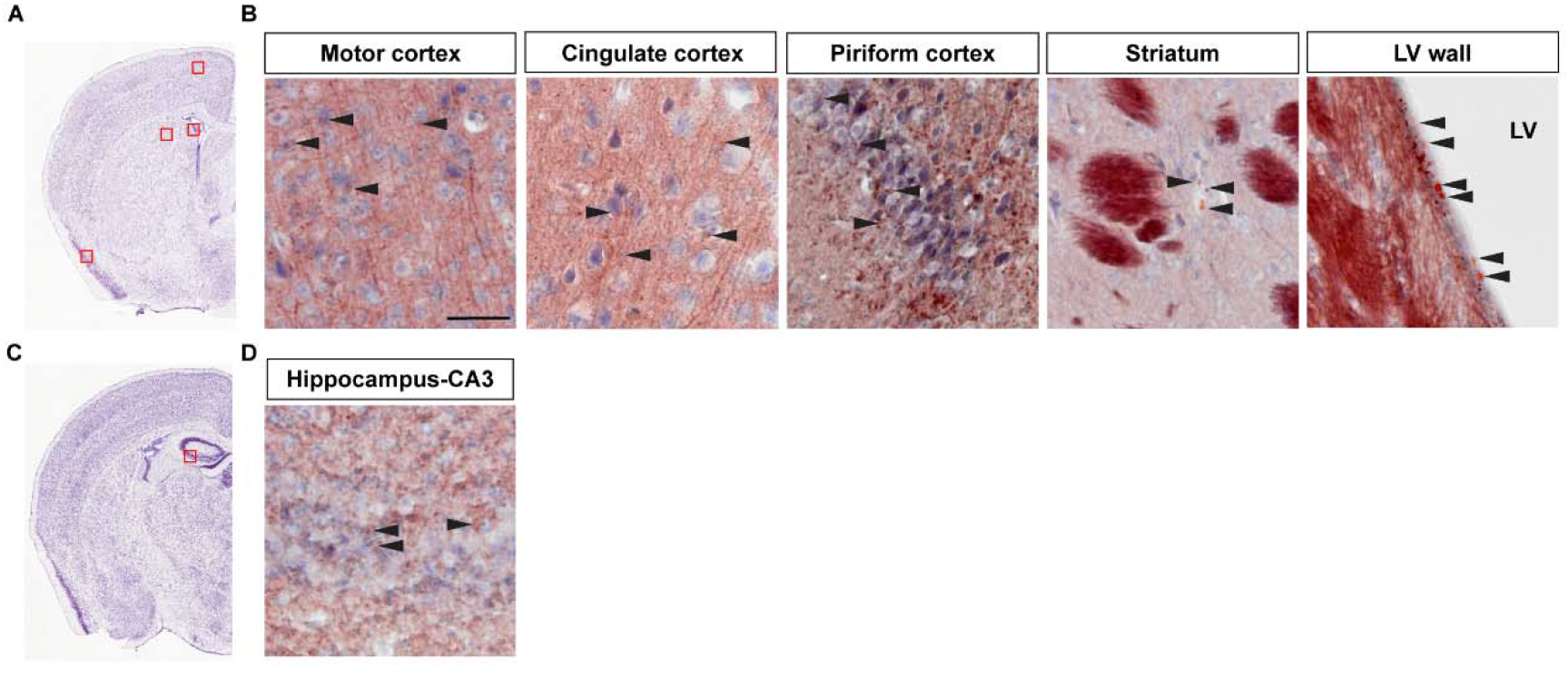
Validation of the ORO staining using aging mouse brain. A 12 month old mouse brain was used to test ORO staining. (A, C) Red boxes in Allen Mouse Brain Atlas images show the brain areas where representative images were taken. (B, D) Arrowheads indicate LDs stained with ORO. Scale bar = 50µm. LV = lateral ventricle.

**Figure 3 – figure supplement 2.**
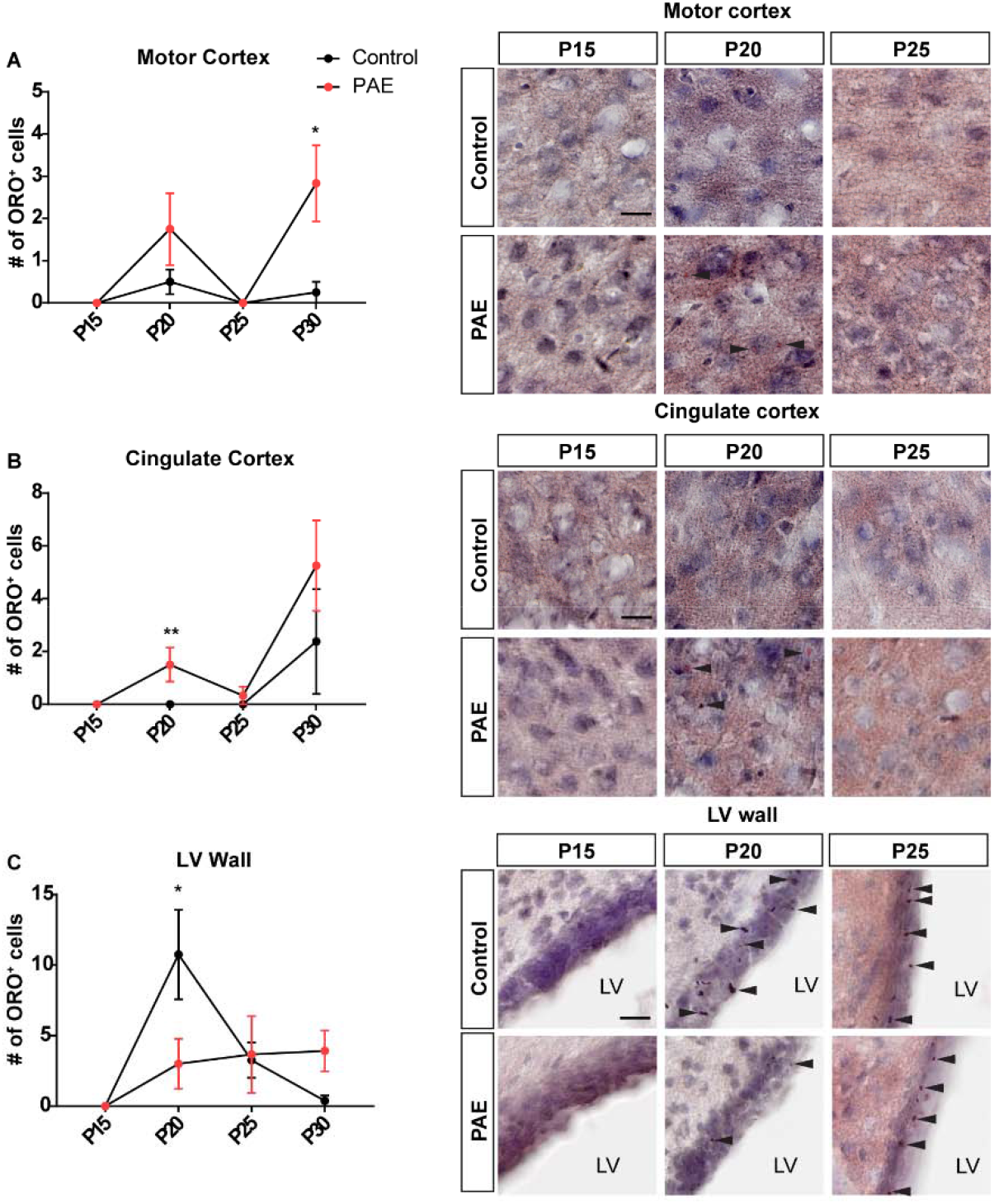
LD accumulation shows dynamic change during postnatal brain development. (A-C) The number of LD accumulating cells is quantified in 200 x 200 µm^2^ area for each brain region and compared between control and PAE mice. At P20, a slight increase of LD accumulating cells in the motor cortex of PAE is observed (p=0.1576). In addition, a significant increase of LD accumulating cells in the cingulate cortex of PAE mice (p=0.0056), whereas a significant increase of LD including cells in the LV wall of control mice (p=0.0226) is observed at P20. At P15 and P25, there is no difference between control and PAE mice in all three brain regions. Two-way ANOVA showed no interaction between the prenatal exposure type and different time points of the observation. Bonferroni’s post hoc test was used to detect statistical significance between control and PAE for each time point. n=4 per group for P15 and P20 time points. n=4 control and n=3 PAE for P25 time point. *p<0.05. Line graphs represent mean ± SEM. P30 data from Figure 3 is added to the graphs for reference. (D) Representative images of ORO staining. Arrowheads indicate LDs. Scale bar = 20µm. LV = lateral ventricle. **Figure 3 – figure supplement 2– source data 1** **Quantification of the number of LD accumulated cells in various brain regions at different time points.**

**Figure 4 – figure supplement 1.**
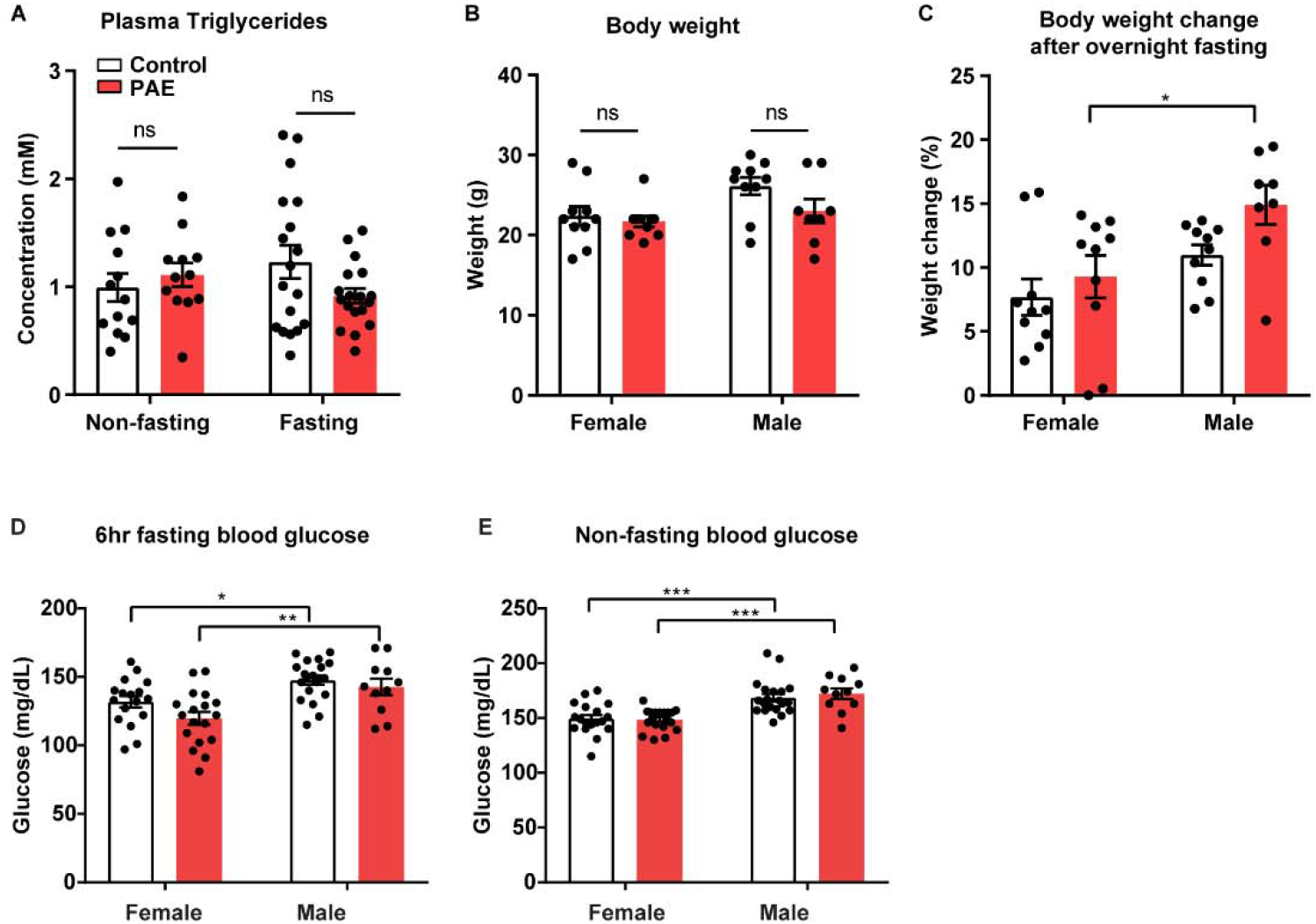
PAE does not affect blood triglyceride, glucose levels, or body weight. (A) No statistical significance is found between control and PAE for plasma triglycerides levels in non-fasted and overnight fasted (16hr) conditions. Although all data from both male and female are used for analysis, no sex dimorphism was observed in either control or PAE by Student’s t-test. Fasting control n=18, Fasting PAE n=18, Non-fasting control n=13, Non-fasting PAE n=12. (B, C) There is no significant difference in body weights between control and PAE mice for both sexes. Body weight after overnight fasting was similar between control and PAE mice. However, in PAE group, male mice lose significantly more weight compared to the female mice after overnight fasting (p=0.0427). No interaction between sex and prenatal exposure type was found by two-way ANOVA. Tukey’s multiple comparisons were used for the post hoc test. Control male = 10, Control female = 10, PAE male = 8, PAE female = 10. (D, E) Two-way ANOVA revealed no interaction between sex and prenatal exposure type. Tukey’s multiple comparison test showed a significant difference between the sexes. There is no significant difference in blood glucose levels between control and PAE mice in both non-fasting and 6 hours fasting conditions. 6 hr fasting: Control female vs Control male (p=0.047), and PAE female vs PAE male (p=0.0074). Non-fasting: Control female vs Control male (p=0.0005), PAE female vs PAE male (p=0.0002). 6hr fasting blood glucose: Control male n=20, Control female n=17, PAE male n=11, PAE female n=18; Non-fasting blood glucose: Control male=20, Control female n=18, PAE male n=11, PAE female n=19. All graphs represent mean ± SEM. Each dot represents an individual animal. *, **, *** p<0.05, 0.01, 0.001. **Figure 4 – figure supplement 1 – source data 1** **Measurements of plasma triglycerides, body weight, change of body weight after overnight fasting, 6hr and non-fasting blood glucose levels.**

**Figure 4 – figure supplement 2.**
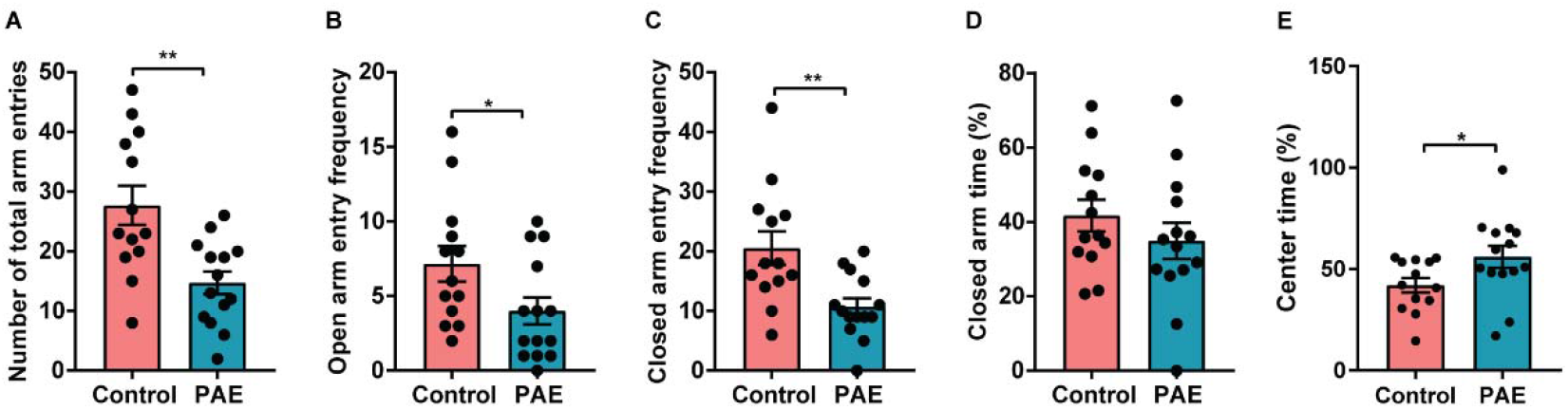
Both numbers of arm entries and center time in EPM are significantly changed in PAE mice compared to control mice. In EPM, the total number of arm entries (open and closed) is significantly decreased in PAE mice compared to control mice (p=0.0018). (B, C) The open arm and closed arm entry frequencies are significantly lower in PAE mice compared to control mice (p=0.0433 and p=0.0038, respectively). (D) No difference in closed arm time is observed between PAE and control mice. PAE mice spend significantly more time in the center of EPM compared to control mice (p=0.0440). Control n=13, PAE n=14. *, ** p<0.05, 0.01. Student’s t-test. All graphs represent mean ± SEM. Each dot represents an individual animal. **Figure 4 – figure supplement 2 – source data 1** **Quantification of the number of total arm entries, open arm entry frequency, closed arm frequency, closed arm time, and center time in EPM test.**

**Figure 4 – figure supplement 3.**
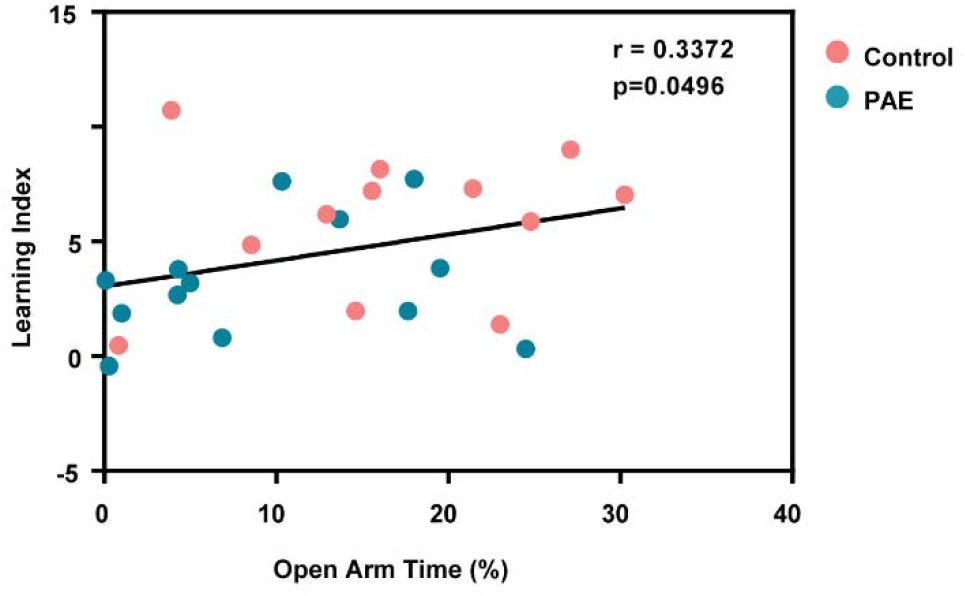
Anxiety phenotype is correlated with poor motor learning behavior. There is a positive correlation between the learning index and open arm time (%) by Pearson’s correlation analysis. Control n=12, PAE n=13. Each dot represents an individual animal. **Figure 4 – figure supplement 3 – source data 1** **Measurements of open arm time and accelerated rotarod learning index.**

**Figure 4 – figure supplement 4.**
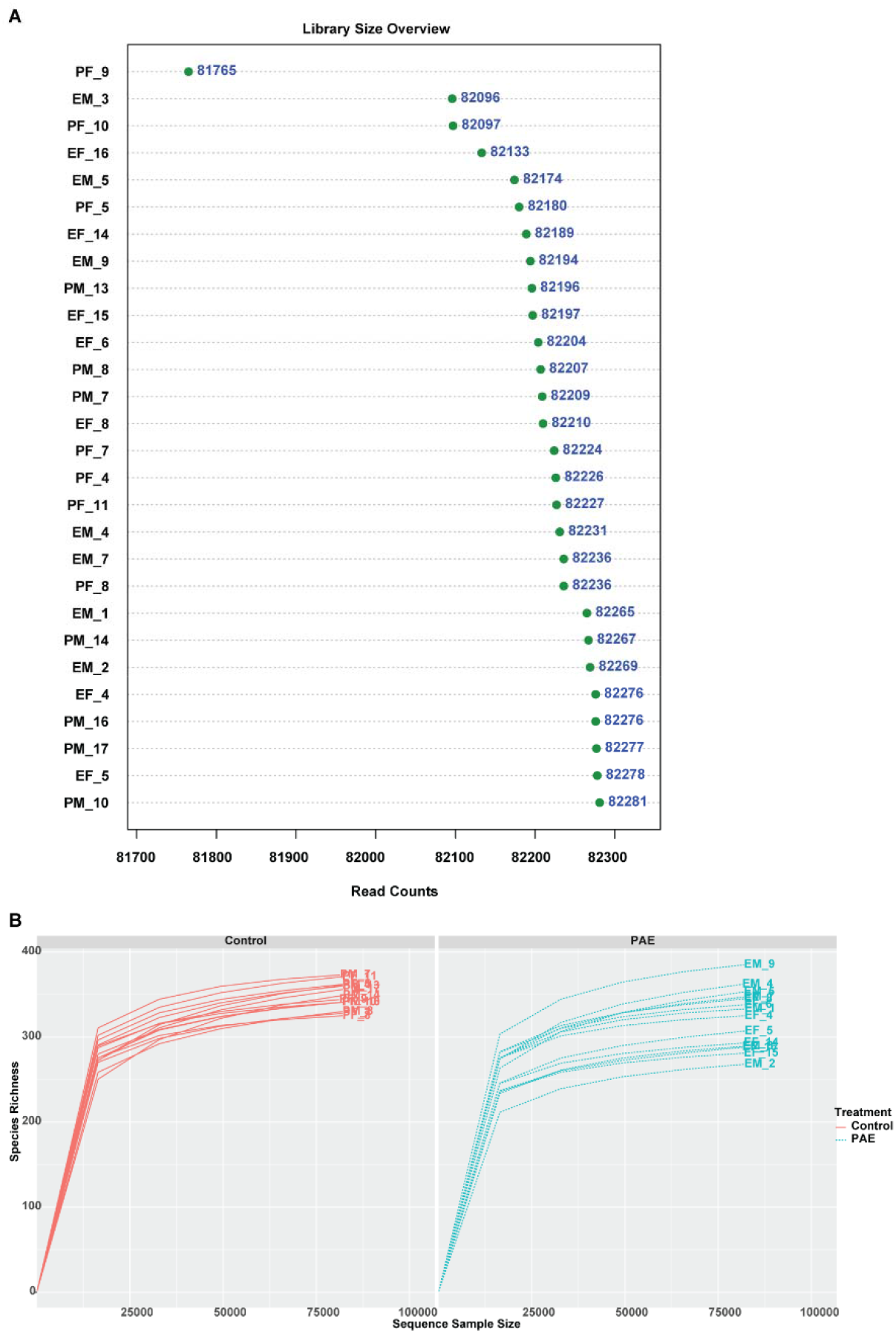
Quality control of 16s rRNA sequencing. (A) Library sizes of samples range from 81765 to 82281 reads. (B) Rarefaction curves show species richness and diversity. Each line represents a sample that is derived from an individual mouse. PF = Control (PBS exposed) female, PM = Control male, EF = PAE female, EM = PAE male.

**Figure 4 – figure supplement 5.**
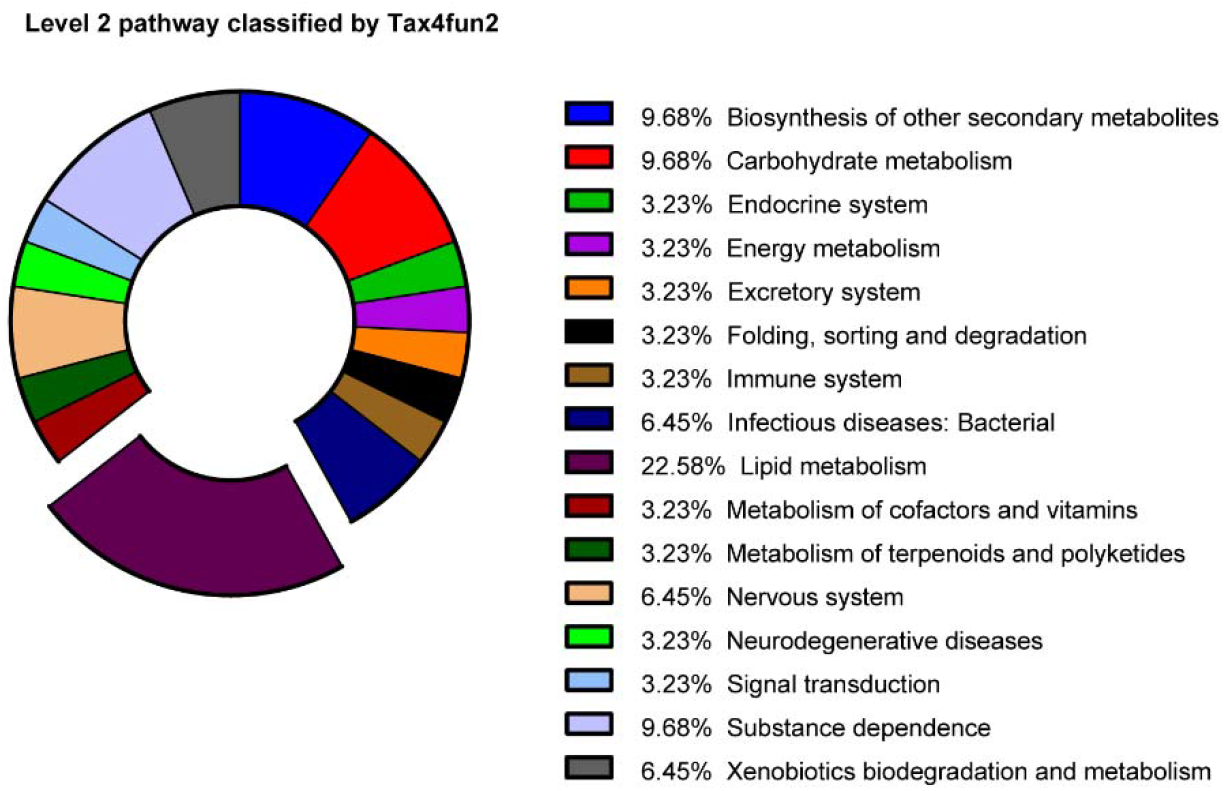
Functional profiles altered by PAE in the microbial community. Pie chart presents the percentages of significantly altered (p<0.05 by Welch’s t-test) pathways in PAE mice compared to the control. Actual percentages of each pathway are indicated in the legend. Lipid metabolism is the profile that shows the largest difference between microbial communities of PAE and control mice.

